# Complete genome of the Medicago anthracnose fungus, *Colletotrichum destructivum*, reveals a mini-chromosome-like region within a core chromosome

**DOI:** 10.1101/2023.12.16.571984

**Authors:** Nicolas Lapalu, Adeline Simon, Antoine Lu, Peter-Louis Plaumann, Joëlle Amselem, Sandrine Pigné, Annie Auger, Christian Koch, Jean-Félix Dallery, Richard J. O’Connell

## Abstract

*Colletotrichum destructivum* (*Cd*) is a phytopathogenic fungus causing significant economic losses on forage legume crops (*Medicago* and *Trifolium* species) worldwide. To gain insights into the genetic basis of fungal virulence and host specificity, we sequenced the genome of an isolate from *M. sativa* using long-read (PacBio) technology. The resulting genome assembly has a total length of 51.7 Mb and comprises 10 core chromosomes and two accessory chromosomes, all of which were sequenced from telomere to telomere. A total of 15,631 gene models were predicted, including genes encoding potentially pathogenicity-related proteins such as candidate secreted effectors (484), secondary metabolism key enzymes (110) and carbohydrate-active enzymes (619). Synteny analysis revealed extensive structural rearrangements in the genome of *Cd* relative to the closely-related Brassicaceae pathogen, *C. higginsianum*. In addition, a 1.2 Mb species-specific region was detected within the largest core chromosome of *Cd* that has all the characteristics of fungal accessory chromosomes (transposon-rich, gene-poor, distinct codon usage), providing evidence for exchange between these two genomic compartments. This region was also unique in having undergone extensive intra-chromosomal segmental duplications. Our findings provide insights into the evolution of accessory regions and possible mechanisms for generating genetic diversity in this asexual fungal pathogen.

**Impact statement:** *Colletotrichum* is a large genus of fungal phytopathogens that cause major economic losses on a wide range of crop plants throughout the world. These pathogens vary widely in their host specificity and may have either broad or narrow host ranges. Here, we report the first complete genome of the alfalfa (*Medicago sativa*) pathogen, *Colletotrichum destructivum*, which will facilitate the genomic analysis of host adaptation and comparison with other members of the Destructivum species complex. We identified a species-specific 1.2 Mb region within chromosome 1 displaying all the hallmarks of fungal accessory chromosomes, which may have arisen through the integration of a mini-chromosome into a core chromosome and could be linked to the pathogenicity of this fungus. We show this region is also a focus for segmental duplications, which may contribute to generating genetic diversity for adaptive evolution. Finally, we report infection by this fungus of the model legume, *Medicago truncatula*, providing a novel pathosystem for studying fungal-plant interactions.

**Data summary:** All RNA-seq data were submitted to the NCBI GEO portal under the GEO accession GSE246592. *C. destructivum* genome assembly and annotation are available under the NCBI BioProject PRJNA1029933 with sequence accessions CP137305-CP137317.

Supplementary data (genomic and annotation files, genome browser) are available from the INRAE BIOGER Bioinformatics platform (https://bioinfo.bioger.inrae.fr/). Transposable Elements consensus sequences are also available from the French national data repository, research.data.gouv.fr with doi 10.57745/TOO1JS.

## Introduction

The ascomycete fungal pathogen *Colletotrichum destructivum*, causes anthracnose disease on lucerne (alfalfa, *Medicago sativa*) and *Trifolium* species and is responsible for significant economic losses on these forage legumes [1, 2]. Despite being isolated most frequently from members of the Fabaceae, *C. destructivum* has occasionally been recorded from genera of the Asteraceae (*Helianthus*, *Crupina*), Poaceae (*Phragmites*) and Polygonaceae (*Rumex*) [3, 4]. It has a worldwide distribution that includes the USA, Canada, Argentina, Italy, Netherlands, Greece, Serbia, Morocco, Saudi Arabia, and Korea. *C. destructivum* is a haploid fungus with no known sexual stage [3]. Previous reports of a sexual stage (*Glomerella glycines*) for soybean isolates of *C. destructivum* [5, 6] were based on incorrect identification of the soybean pathogen, which was recently shown to be *C. sojae* [7].

Over the last decade, the application of multi-locus molecular phylogeny approaches has revealed that *C. destructivum* belongs to the Destructivum species complex, which contains 17 accepted taxa [3, 8]. All these plant pathogenic species show distinct host preferences, spanning phylogenetically diverse botanical families. An increasing number of species in the Destructivum complex have now been genome sequenced, namely *C. higginsianum* [9, 10], *C. tanaceti* [11], *C. lentis* [12] and *C. shisoi* [8], which cause disease on Brassicaceae, *Tanacetum* (Asteraceae), *Lens* (Fabaceae) and *Perilla* (Lamiaceae), respectively. The clade therefore provides excellent opportunities for comparative genomic studies on the genetic determinants of host adaptation.

The availability of complete genome sequences is crucial not only for the analysis of large gene clusters, such as secondary metabolism biosynthetic gene clusters, but also for understanding fungal genome evolution. Complete or near-complete genome sequences have enabled the structure and dynamics of accessory mini-chromosomes to be analyzed in several *Colletotrichum* species [9, 13, 14]. The importance of mini-chromosomes for virulence on plant hosts has been demonstrated in several fungal pathogens including *Fusarium oxysporum* f.sp. *lycopersici* [15], *Magnaporthe oryzae* [16], *C. lentis* [12] and *C. higginsianum* [17].

Here, we present the complete genome sequence and gene annotation of *C. destructivum* strain LARS 709, hereafter called *Cd*709, based on long-read sequencing with PacBio Single Molecule, Real-Time (SMRT) Sequel technology. The resulting high-quality chromosome-level assembly allowed us to perform comparative genomics with the close sister species, *C. higginisianum*, highlighting gene content specificity and extensive genomic rearrangements. In particular, the genome showed evidence of multiple segmental duplications, as well as the likely integration of a mini-chromosome into one core chromosome. Although the origin of this integrated region remains to be determined, it displays all the hallmarks of fungal mini-chromosomes. We also show for the first time that *C. destructivum* is pathogenic, and completes its life-cycle, on the model plant *Medicago truncatula*, providing a new tractable pathosystem in which both partners have been genome-sequenced.

## Materials and Methods

### Fungal and plant materials

The *C. destructivum* strains used in this study were originally isolated from *M. sativa* in Saudi Arabia (CBS 520.97, LARS 709) and Morocco (CBS 511.97, LARS 202) [2], and are hereafter called *Cd*709 and *Cd*202. The *C. higginsianum* strains used for comparative genome and chromosome analyses were IMI 349063A and MAFF 305635 [10, 17, 18], hereafter called *Ch*63 and *Ch*35, respectively. The fungi were cultured as described previously [18].

Seeds of nine *M. truncatula* accessions (Table S1) were provided by the INRAE Centre de Ressources Biologiques Medicago truncatula (UMR 1097, Montpellier, France), while *M. sativa* seeds were purchased from Germ’line SAS (France). *M. truncatula* seeds were first abraded with sandpaper and imbibed with water for 1 h before sowing in seed compost (Floragard Vertriebs-GmbH, Oldenburg, Germany), while *M. sativa* seeds were sown directly in the same compost. All plants were grown in a controlled environment chamber (23°C day, 21°C night, 12-h photoperiod, PPFR 110 μmol·m^-2^·s^-1^).

### Infection assays and microscopy

To test the susceptibility of *M. truncatula* accessions to *Cd*709, intact plants (17-days-old) were inoculated by first immersing the above-ground parts in a solution of 0.01 % (v/v) Silwet to wet the leaves, then by immersion in a suspension of *C. destructivum* spores (2 x 10^6^ ml^-1^). The inoculated plants were incubated in a humid box inside a controlled environment chamber (25°C, 12-h photoperiod, PPFR 40 μmol m^-2^ s^-1^). For microscopic examination, pieces of infected tissues were cleared with a 1:3 mixture of chloroform:ethanol for 1h, then with lactophenol for 30 min, before mounting on a microscope slide in 70 % glycerol and imaging with a Leica DM5500 light microscope. Symptoms were recorded at 4 dpi.

### Pulsed-field gel electrophoresis (PFGE) and Southern blotting

The plugs containing the conidial protoplasts for PFGE were prepared as previously described [17]. Pulsed-field gel electrophoresis (Bio-rad CHEF-DR II system) was performed using the following conditions: Runtime 260 hours; Switch time 1200 s to 4800 s; 1.5 V / cm; 0.75 x TBE at 8°C. Yeast chromosomal DNA served as size marker (BioRad; 200 kb – 2 Mb).

Southern blotting was conducted using standard protocols [19]. A digoxigenin labeled probe was generated by PCR following the manufacturer’s instructions (PCR DIG Probe Synthesis Kit,Roche). The 993 bp probe (*Cd*709 chr1, position 6,711,095 to 6,712,088) was specific to mini-chromosome-like sequences at the right arm of chromosome 1 in *Cd*709. Hybridization was performed in DIG Easy Hyb buffer at 42°C overnight. The membrane was then extensively washed with low and high stringency buffers and subsequently blocked with buffer B2 (1% Blocking powder [Roche] in buffer B1 [100 mM Maleic acid, 150 mM NaCl, pH 7.5]). The blocking solution was then replaced with antibody solution (buffer B2 containing DIG-antibody 1:26,000 (Roche)). The membrane was washed with buffer B1 containing 0.3% Tween20. The membrane was subsequently equilibrated in buffer B3 (100 mM Tris pH 9.5, 100 mM NaCl, 50 mM MgCl_2_) and developed with chemiluminescence (CDP-Star, Roche).

### Genome data, assembly, rearrangements and duplications

The genomic DNA of *Cd*709 was used to prepare a size-selected library (20kb) prior to sequencing with a PacBio Sequel sequencer (kit 2.1, Keygene N.V., Wageningen, The Netherlands) on two SMRT cells, yielding raw data with approximately 224 X genome coverage (1.474.759 reads, N50 10.837 bp). Genome assemblies were generated from several runs of the Hierarchical Genome-Assembly Process version 4 (HGAP4) and Canu [20] assemblers. The draft genome was polished with the Arrow algorithm and the completeness of the assembly was evaluated with BUSCO using the Ascomycota gene set as evidence [21]. Telomeres were validated by the presence of at least three repeats of the TTAGGG/CCCTAA motif at the end of assembled contigs [22]. The polished assembly was aligned with nucmer against the *Ch*63 and *Ch*35-RFP genomes to visualize chromosome rearrangements. SDDetector [9] was used to detect segmental duplications in combination with Bedtools and BWA-MEM for validation. The *Cd*709 mitochondrial genome was assembled with Organelle_PBA [23] (Table S2).

### Transcriptome data and analysis

RNA sequencing was performed on samples of mRNA from undifferentiated mycelium grown axenically and two different stages of plant infection, 48 and 72 h after inoculation, corresponding to the biotrophic and necrotrophic phase, respectively. Mycelium was grown for three days in potato dextrose liquid medium (PDB, Difco) at 25°C with shaking (150 rpm) and harvested by filtration. Seedlings of *M. sativa* (eight days old) were inoculated by placing a droplet (10 μl) of *Cd*709 spore suspension (7 x 10^5^ spores/ml) onto the surface of each cotyledon and the plants were then incubated as described for *M. truncatula*. Discs of infected cotyledon tissue were harvested using a cork borer (4 mm diameter). After grinding the tissues in liquid nitrogen, total RNA was extracted using the RNeasy plant mini kit (Qiagen). Libraries were then prepared from each sample type using the TruSeq Paired-end Stranded mRNA Kit and sequenced (100 bp reads) using a HiSeq4000 sequencing platform (IntegraGen Genomics, Evry, France). RNA-Seq paired reads were cleaned and trimmed using Trimmomatic [24] and then mapped to the genome assembly of *Cd*709 using STAR [25]. A genome-guided transcript assembly was obtained from mappings with StingTie v1.3.4. Assembled raw transcripts were then filtered based on the TPM distribution per transcript per library.

### Genome annotation

Transposable elements (TE) were searched in the *C. destructivum* genome sequence using the REPET package [26, 27]. Consensus sequences identified with the TEdenovo pipeline were classified using the PASTEC tool [28], based on the Wicker hierarchical TE classification system [29], and then manually filtered and corrected. The resulting library of consensus sequences was used to annotate TE copies in the whole genome using the TEannot pipeline.

Protein-coding genes were annotated using the Eugene [30] and FunGAP [31] tools. Predicted genes were filtered out when 10% of their CDS overlapped a Transposable Element predicted by the REPET package. Filtered predicted genes from Eugene and FunGAP were clustered together based on their CDS coordinates (overlap of one base required) with no strand consideration. The Annotation Edit Distance (AED) [32] was computed with transcript and protein evidence for each transcript and the predicted model with the best score was retained at each locus. Mitochondrial genomes were annotated with MFannot [33] and MITOS2 [34]. Results were manually inspected and in case of divergence between the predictions, the longer gene model was retained.

The synteny between *C. destructivum* and *C. higginsianum* proteomes was analysed with SynChro [35] which detects ortholog proteins with Reciprocal Best Hit (RBH), based on 40% similarity and a length ratio of 1.3. Colinear orthologs were then grouped in syntenic blocks, according to a delta threshold = 1 (very stringent mode). Non-syntenic blocks were extracted when five or more consecutive non-syntenic genes were found. Proteome similarities with other *Colletotrichum* spp. were performed with Blast 2.2.28+ and the results filtered with a cut-off of 30% identity and 50% query coverage. Proteome synteny and associated figures were obtained using Clinker [36].

### Functional annotation of predicted genes

Functional annotations of genes obtained using Interproscan 5.0 [37] and Blastp (e-value <1e-5) [38] against the NCBI nr databank (September 2019) were then used to perform Gene Ontology [39] annotation with Blast2GO [40]. Carbohydrate active enzymes (CAZymes) were annotated with dbCAN2 [41] launching HMMER, Diamond and Hotpep against dedicated databases. Genes were considered as CAZymes when at least 2 of the three tools provided a positive annotation.

Genes encoding potential secreted proteins were predicted with a combination of SignalP v4.1 [42], TargetP v1.1 [43] and TMHMM v2.0 [44] results. The secretome was defined as the union of SignalP and TargetP results and then intersected with TMHMM results (0 or only 1 transmembrane domain).

Proteins smaller than 300 amino acids were then extracted and considered as Small Secreted Proteins (SPPs). In parallel, EffectorP v2.0 [45] was applied to the predicted secretome to identify putative effector proteins. Finally, the intersection of EffectorP and SPPs results was retained to establish a list of potential effectors.

To detect secondary metabolism biosynthetic gene clusters (BGCs), predicted genes were submitted to antiSMASH (Antibiotics and Secondary Metabolite Analysis Shell) v5 [46]. Only core biosynthetic genes (commonly known as secondary metabolism key genes, SMKGs) were considered for further analysis. Presence/absence patterns of SMKGs were based on reciprocal best hits with *Ch*63 and *Ch*35, and then manually inspected. Among the newly predicted secondary metabolism key genes (SMKGs), those encoding polyketide synthases (PKS) and non-ribosomal peptide synthases (NRPS) were checked for the presence of the minimal expected set of enzymatic domains, namely KS and AT domains for PKS, and A and PCP domains for NRPS. Terpene synthases and dimethylallyltryptophan synthase (DMATS) genes were manually inspected and retained if they had RNA-seq or protein support. Those *Cd*709 genes not predicted as SMKGs by antiSMASH, but orthologous to a *C. higginsianum* SMKG were also included. For example, antiSMASH failed to annotate six terpene synthase (TS) that are present in both species.

### Codon usage analysis

Codon usage was computed for predicted gene coding sequences (CDS) on each chromosome or region using the EMBOSS tool ‘cusp’. The resulting codon usage matrix (i.e. the fraction of each codon in a given amino acid) was subjected to Fisher’s exact tests (with a Bonferroni correction for multiple testing) to address the statistical significance of differences between the core and mini-chromosomes. The matrix was also subjected to a Principal Component Analysis (PCA) and the results were projected onto the first two principal components. To analyse the GC percentage of the three letters of each codon, the ‘cusp’ tool was run individually on each CDS of each chromosome or region and the results were represented as density plots. The corresponding figures were generated using R (v. 4.0.5) and the libraries ggplot2 (v. 3.3.3), cowplot (v.1.1.1) and ggbeeswarm (v. 0.6.0), all available from the CRAN repository (https://cran.r-project.org/).

## Results

### A novel Colletotrichum destructivum - Medicago truncatula pathosystem

The cell biology of infection of *M. sativa* by *C. destructivum* isolate 709 (*Cd*709) was previously described [2]. Here, we report infection of the model plant *M. truncatula* (barrel medic) by this species. Five out of the nine tested *M. truncatula* accessions, including the genome-sequenced accession ESP074-A [47], were found to be susceptible to *C. destructivum* in two independent infection assays (Fig. 1, Table S1). At 4 days post inoculation (dpi), necrotic water-soaked lesions were visible on the trifoliate leaves of the susceptible accessions (Fig. 1). In contrast, the leaves of resistant accessions presented only small necrotic flecks or no visible symptoms. The genome-sequenced accession R108-C3, which is widely used for *M. truncatula* functional genomics [48], was resistant to *C. destructivum* in these infection assays.

**Figure 1:**
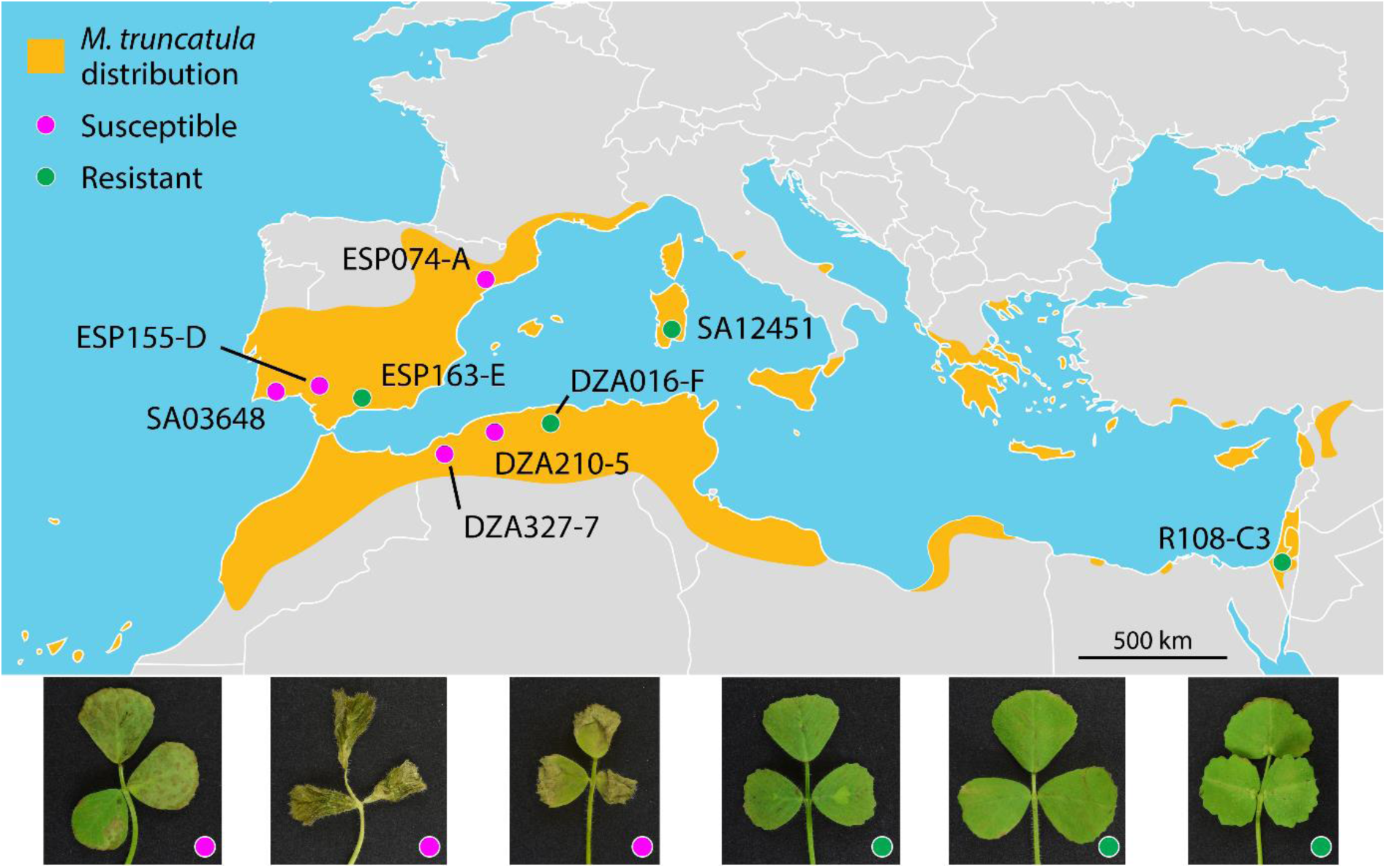
*Medicago truncatula* accessions used in this study and their infection phenotypes with *Colletotrichum destructivum* LARS 709. **Upper panel:** Geographical distribution of *M. truncatula* in the Mediterranean area according to GBIF (2019) and collection locations of the nine ecotypes used in this study. **Lower panel:** Symptoms produced on the trifoliate leaves of six *M. truncatula* accessions at 4 days post inoculation with spore suspension of *C. destructivum* LARS 709. Leaves of the susceptible accession DZA210-5 showed large necrotic lesions, while those of DZA327-7 and ESP155-D were completely necrotic. Leaves of the resistant accessions ESP163-E, DZA016-F and R108-C3 showed small necrotic flecks or no visible symptoms. Note that R108-C3 is considered to be *M. truncatula* ssp. *tricycla*.

On cotyledons of the susceptible *M. truncatula* accession ESP155-D, *Cd*709 spores germinated to form melanized appressoria, which by 48 hpi had penetrated host epidermal cells to form bulbous, intracellular biotrophic hyphae that were confined to the first infected cell (Fig. 2a). Thinner necrotrophic hyphae started to emerge from the tips of the biotrophic hyphae at 60 hpi (Fig. 2b), and after 72 hpi the fungus had completed its asexual cycle by producing sporulating structures (acervuli) on the surface of the dead tissues (Fig. 2c). On cotyledons of the resistant accession ESP163-E, appressoria formed abundantly on the leaf surface but penetrated host epidermal cells very infrequently (Fig. 2d, e). Groups of dead epidermal cells underlying the appressoria appeared yellow-brown in colour and had granulated contents, suggesting they had undergone a hypersensitive cell death response. Rarely, small hyphae were visible in epidermal cells beneath appressoria but they developed only a short distance into the dead cells and most remained smaller than the appressorium. Acervuli were never observed on plants of accession ESP163-E.

**Figure 2:**
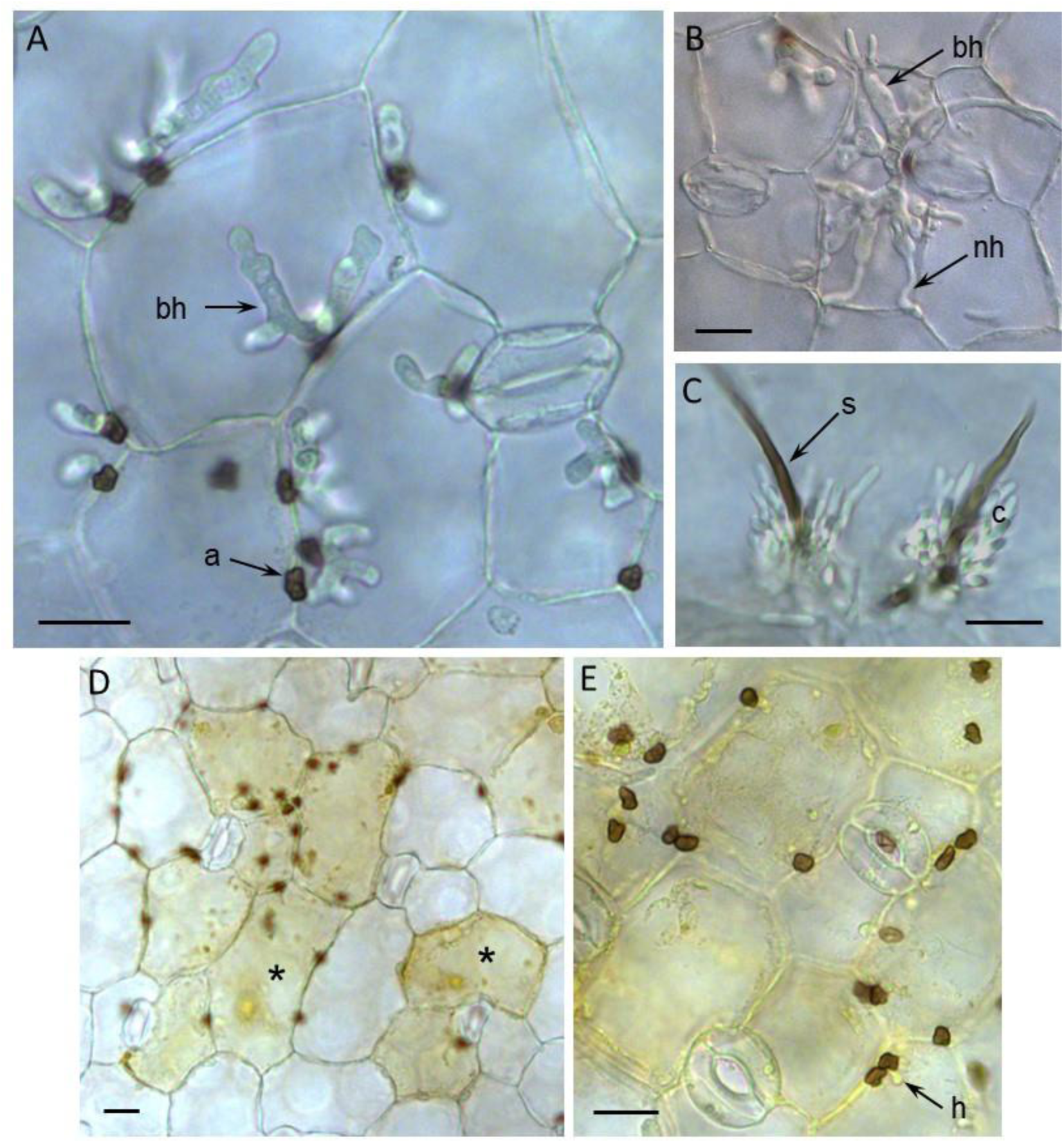
Microscopic analysis of *Colletotrichum destructivum* LARS 709 infecting cotyledon tissues of *Medicago truncatula*. **(A-C)** Susceptible accession ESP155-D. At 48 hpi **(A)**, melanized appressoria (a) had formed on the plant surface and penetrated epidermal cells to form bulbous biotrophic hyphae (bh). At 60 hpi **(B)**, thin necrotrophic hyphae (nh) developed from the tips of biotrophic hyphae. At 72 hpi **(C)**, acervuli erupted from the plant surface, consisting of a melanized, hair-like seta (s) and a mass of conidia (c). **(D,E)** Resistant accession ESP163-E. At 72 hpi, few appressoria had penetrated cotyledon epidermal cells, and groups of cells underlying the appressoria were pigmented yellowish brown with granular contents (*). Any hyphae (h) visible inside epidermal cells were typically smaller than the appressorium. Scale bars = 20 μm.

### Genome assembly and structural annotation

Long-read data allowed us to generate a complete genome assembly for *Cd*709, with a total length of 51.75 Mb in which all 12 chromosomes were sequenced from telomere to telomere (Fig. 3), together with the circular mitochondrial genome (34 kb). Annotation of transposable elements revealed a total of 49 consensus sequences, representing all the possible TEs in the *Cd*709 genome. Classification of the TEs (Table S3) showed that the genome contains 18 different families of retrotransposons, including eleven LTR (Long Terminal Repeats) and seven LINE (long interspersed nuclear element), 28 DNA transposons, including 25 TIR (terminal inverted repeat), one helitron and two MITE (Miniature Inverted-Repeat Transposable Elements), as well as three unclassified repeated elements. The library of 49 consensus sequences was then used to annotate TE copies in the *Cd*709 genome. Overall, TEs covered 6.2 % of the genome assembly by length. The Class I LTR Gypsy superfamily was the most abundant in terms of coverage and number of copies, whereas the Class I TIR Tc1-Mariner was the most abundant in terms of full-length copies. Two Gypsy transposons (R172 and G87) resemble the most abundant TE family in *C. higginsianum*, namely the LTR transposon family RLX_R119 [9]. Looking at the distribution of TE families along the chromosomes, we found that the telomeres of all twelve *C. destructivum* chromosomes were associated with a single copy of a TE belonging to the helitron family (G103).

**Figure 3:**
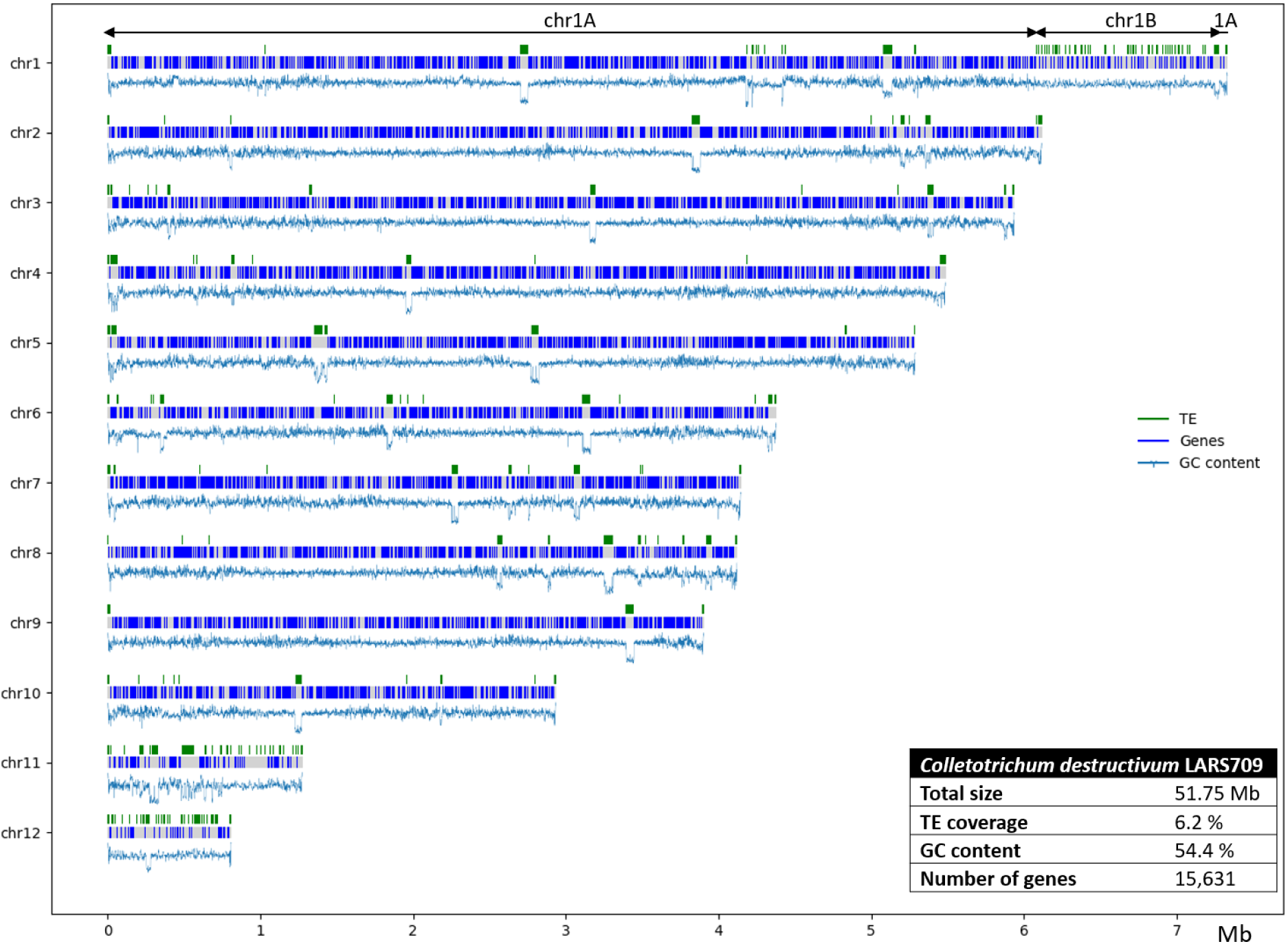
Schematic representation of the 12 chromosomes of *Colletotrichum destructivum* isolate 709. The distribution of genes and transposable elements (TE) across each chromosome are shown together with the corresponding genome statistics (inset table).

To annotate the protein-coding genes, a genome-guided assembly of RNA-Seq reads provided 16,122, 13,901 and 15,081 transcripts for axenic mycelium, 48 hpi and 72 hpi libraries, respectively (Table S4), with 1.88 TPM, 9.38 TPM and 4.90 TPM as minimum expression levels, respectively (Fig. S1). Assembled transcripts were then used to predict gene models in conjunction with *Colletotrichum* and Ascomycota protein databanks. The results of EuGene and FunGap were combined and filtered to generate the *Cd*709 gene set comprising 15,631 complete gene models, of which 11,853 had transcript support and 15,172 resembled Ascomycota predicted proteins. Features of the gene annotation are summarized in Table S5. The completeness of this annotation was confirmed by comparison to the BUSCO Ascomycota set (1,315 genes), with 1,309 complete genes predicted and only one missing. Functional annotation assigned InterPro entries to 10,298 genes, among which 7,475 had at least one GO term and 1,105 were potential enzymes (annotated with an Enzyme Code). Based on Blast2GO descriptions, 12,192 predicted genes (78%) had a predicted function, i.e. a description other than “hypothetical protein” (Table S6 tab ‘All’). The mitochondrial genome of *Cd*709 was annotated with 29 tRNAs, 2 rRNAs (small and long subunit) and 21 genes.

### Plant interaction-related genes

A total of 619 *Cd*709 genes were annotated to encode CAZymes, among which 410 were assigned to the Glycoside Hydrolase (GH), Carbohydrate Esterase (CE) and Polysaccharide Lyase (PL) CAZyme classes (Table S6 tab ‘CAZyme’). The proportion of genes in each CAZyme class closely resembled that previously found in *Ch*63 [49], and 98% (400/410) of *Cd*709 CAZyme genes were also detected in the *Ch63* genome. *In silico* analysis of the *Cd*709 secretome revealed a total of 2,608 potential extracellular secreted proteins, including 1,118 small proteins (<300 amino acids). Among these, 484 genes were retained as putative effectors because they were also present among 508 genes identified by EffectorP. Comparing these to the effector repertoire of *Ch*63, a total of 127 putative effectors (26.2%) were unique to *Cd*709, having no Reciprocal Best Blast Hit in *Ch*63 (Table S6 tab ‘Predicted effectors’). A total of 110 secondary metabolism key genes (SMKGs) were detected in the *Cd709* genome using the fungal version of antiSMASH and were manually curated. These *C. destructivum* SMKGs were compared to the 105 *C. higginsianum* SMKGs [9]. Overall, 78 % (94 out of 120) of the SMKGs were present in both species (Table S6 tab ‘Secondary metabolism’, Fig. S2). A total of 17 *C. destructivum* SMKGs, distributed over eight BGCs, were not detected in *C. higginsianum*.

### Chromosome structure comparison

Complete chromosome-level assemblies are available for two different *C. higginsianum* strains, namely IMI 349063A (*Ch*63) [9] and MAFF 305635-RFP (*Ch*35-RFP), a transformant of MAFF 305635 (*Ch*35) expressing red fluorescent protein which lacks both mini-chromosomes 11 and 12 [10, 17]. The genetic proximity of *C. destructivum* and *C. higginsianum* allowed us to align assemblies to observe chromosome structural variations. This generated 38 Mb of *C. destructivum* alignments (>10 kb) with each *C. higginsianum* strain, ranging from 88 to 96.7% identity. Thus, *C. destructivum* shared approximately 73.6 % of its total genome length with *C. higginsianum*. At the chromosome scale, alignments revealed that five chromosomes of *C. destructivum* (chr1, 2, 3, 5 and 9) were not involved in any large rearrangements, five others (chr4, 6, 7, 8 and 10) showed inter-chromosomal rearrangements, while the two mini-chromosomes (chr11 and 12) lacked large regions of conserved sequences and appear to be species specific (Fig. 4A).

**Figure 4:**
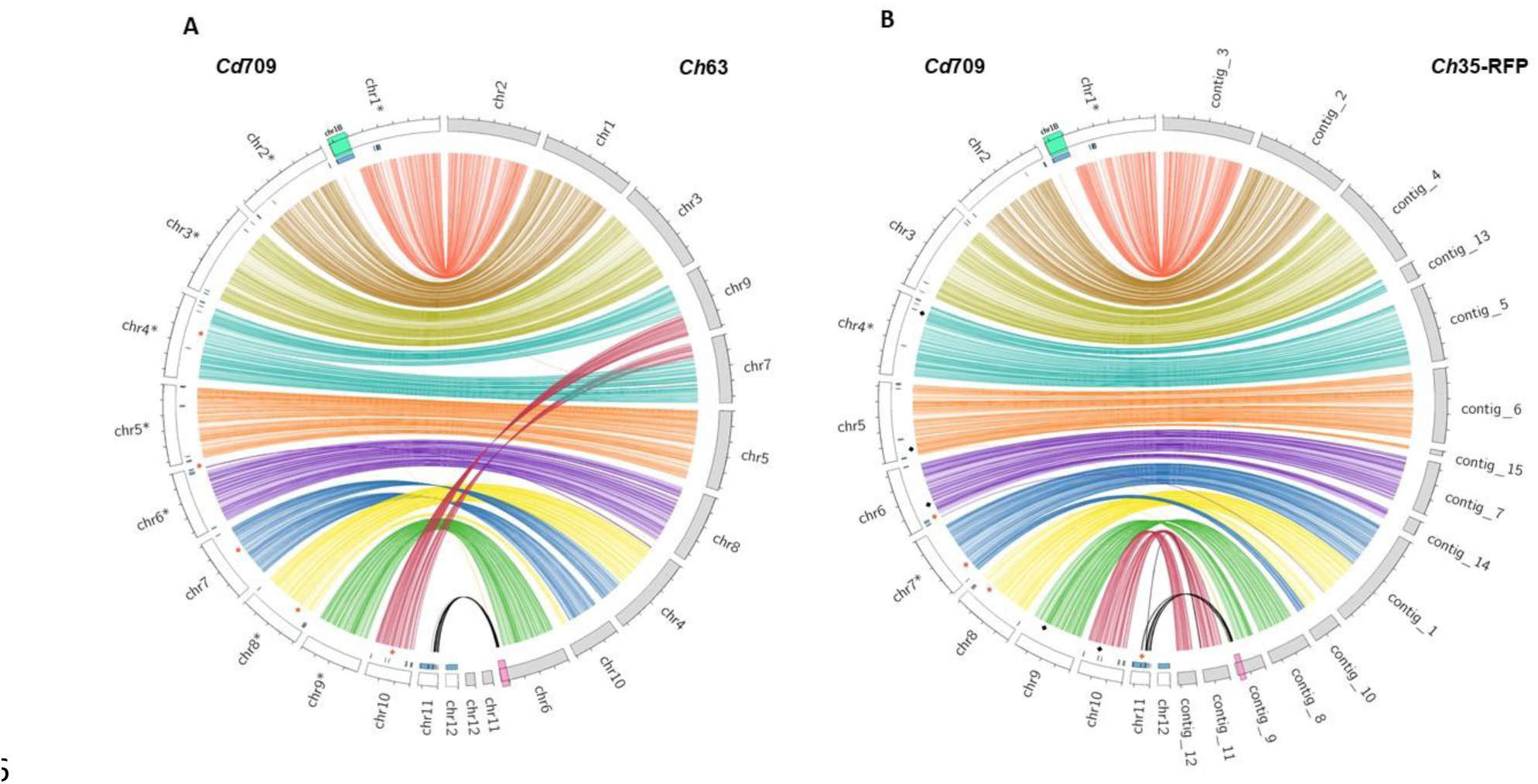
Whole-genome alignments between *Colletotrichum destructivum* LARS 709 (*Cd*709) and two *Colletotrichum higginsianum* strains. Chromosomes of *Cd*709 (white bars) were aligned with **(A)** the chromosomes of *C. higginsianum* IMI 349063 (*Ch*63, grey bars) or **(B)** the contigs of *C. higginsianum* MAFF 304535-RFP (*Ch*35-RFP, grey bars). Syntenic regions (length >10 kb and percent identity > 88%) were linked together using coloured arcs specific for each chromosome in the *Cd*709 genome assembly. Red diamonds indicate interchromosomal rearrangements. Black diamonds indicate chromosome breakpoints associated with separate contigs in the *Ch*35-RFP assembly only. The blue track indicates gene blocks that are unique to *Cd*709. Note that region chr1B of *Cd*709 (highlighted in green) has no alignments in either of the *C. higginsianum* isolates. The black arcs linking chr11 of *Cd*709 to the 3’ end of chr6/contig_9 in *C. higginsianum* (highlighted in pink) indicate regions with strong sequence similarity (percent identity > 88%) that are smaller than 10 Kb. Asterisks indicate where chromosome sequences were reverse-complemented for better visualization. Tick mark spacing = 1 Mb

One rearrangement involved chr7 and chr8 of *Cd*709 resulting in chr4 and chr10 of *Ch*63. The break-points in chr7 and chr8 were associated with TEs in *Cd*709 (Fig. S3 A and B). A similar rearrangement was found relative to *Ch*35-RFP, albeit with different break-points in both species that were not associated with TEs (Fig. S3 F and G). A second rearrangement involved chr4 and chr10 of *Cd*709 such that their left and right arms result in chr9 and chr7 of *Ch*63, respectively (Fig. S3 C and D). Interestingly this rearrangement was not found relative to *Ch*35-RFP, suggesting that it is specific to particular *C. higginsianum* strains, as was noted previously [10]. A third inter-chromosomal rearrangement concerned 121 kb at the 5’ extremity of *Cd*709 chr6 coming from chr4 and contig_1 of *Ch*63 and *Ch*35-RFP, respectively. In *C. destructivum*, this break-point is surrounded by TEs and non-syntenic regions (Fig. S3 E). Remarkably, a specific rearrangement of 42 kb between chr11 of *Cd*709 and contig 11 of *Ch*35-RFP (Fig. S3 H) corresponds to a region that is absent from the *Ch*63 genome assembly and which encodes highly variable effectors (having ≤ 90% alignment coverage) and secondary metabolism-related proteins [10]. In addition, several short stretches (2 to 5 kb in length) from chr11 of *Cd*709 were present at the extremities of chromosome 6 in *Ch*63 and the corresponding region of *Ch*35-RFP (contig_9) (Fig. 4).

A notable feature of the *C. destructivum* genome assembly is the unusually large size of chr1 (7.3 Mb), which is 0.9 Mb longer than the largest chromosome in *C. higginsianum* (6.4 Mb). Genome alignments highlighted a near-complete synteny between chr1 of *Cd*709 and chr2 of *Ch*63 except for a 1.2 Mb subtelomeric region (coordinates chr1:6076875-7282542), for which no similarity was found in *C. higginsianum* (Fig. 4). Synteny between the genes of *Cd*709 and those of *Ch*63 was investigated using SynChro. With stringent settings, 400 syntenic blocks were identified based on 12,135 Reciprocal Best Hits. A total of 1,083 genes were found in 47 non-syntenic blocks composed of at least five consecutive *Cd*709-specific genes (Tables S7 & S8). The largest non-syntenic block, corresponding to the 1.2 Mb region specific to Cd709 on chr1, contained 305 genes. Mini-chromosome chr12 contained one non-syntenic block of 170 genes, while chr11 was divided into seven non-syntenic blocks, the largest containing 106 genes. Although only 356/1,083 genes inside non-syntenic blocks could be annotated with a GO term, GO enrichment tests revealed that the *Cd*709-specific genes were enriched in protein kinases, protein phosphorylation activity and secondary metabolism process (Table S9). Likewise, effector genes were found to be enriched in non-syntenic blocks whereas CAZymes were depleted (Table S10).

Validation of the 1.2 Mb non-syntenic region in *C. destructivum* chromosome 1

To verify the large non-syntenic region identified within chr1, we first checked for potential errors in the sequence assembly of this region by manually inspecting long reads spanning the two junctions (Fig. S4). Secondly, to obtain an assembly-independent validation, pulsed-field gel electrophoresis (PFGE) and a Southern hybridization were performed (Fig. 5A, B). A 993 nt probe (coordinates chr1: 6,711,095 to 6,712,088) was designed within the 1Mb non-syntenic region to target a unique locus that avoided TEs (Fig. 5B). This probe is 83.5% identical to the gene CH63R_14488 located on chromosome 11 of *Ch*63 that was used as a hybridization control.

**Figure 5:**
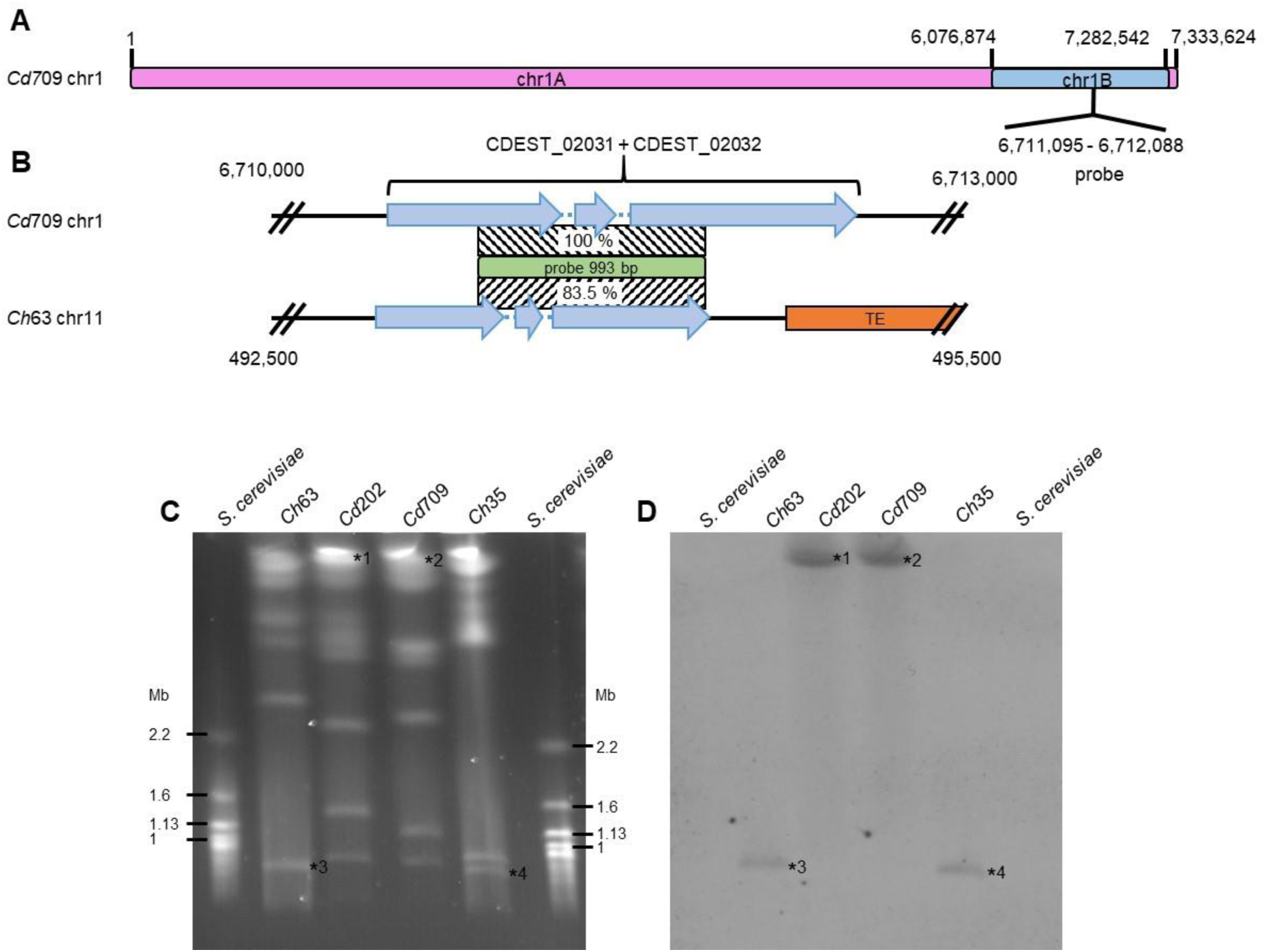
Chromosome 1 of *Colletotrichum destructivum* has a bipartite structure. **(A)** Scheme of the structure of *Cd*709 chromosome 1. The probe is specific to the mini-chromosome-like part of the chromosome (chr1B). **(B)** Detailed scheme of the regions targeted by the 993 bp DIG-labelled probe in *Cd*709 and in *Ch*63 (chr11: 493,380 to 494,373). Patterned boxes indicate sequence identity of the target regions to the probe. **(C)** Pulsed-field gel electrophoresis of chromosomal DNA from *C. destructivum* isolates LARS 202 (*Cd*202) and LARS 709 (*Cd*709) compared to *C. higginsianum* isolates IMI349063 (*Ch*63) and MAFF305635 (*Ch*35). **(D)** Southern hybridisation. Numerals 1 to 4 indicate signals corresponding to chromosomes displayed in **(C)**.

Chromosomes of two *C. destructivum* isolates (*Cd*709 and *Cd*202) and two *C. higginsianum* isolates (*Ch*63 and *Ch*35) were separated by PFGE and analysed by Southern hybridization (Fig. 5C, D). For both *C. destructivum* isolates, the probe hybridized to molecules with high molecular weight that could correspond to the largest chromosome, consistent with a location on chr1 (Fig. 5C, D). The high molecular weight signals were absent in the *C. higginsianum* blots, and instead hybridization signals were detected at a position corresponding to mini-chromosome 11, although these were weak, as expected for a probe with only 83.5% identity to the target. Overall, our findings validate that a non-syntenic region is embedded within chr1 of *C. destructivum*. Hereafter, we refer to the syntenic and non-syntenic portions as chr1A and chr1B, respectively, and their distinct properties were explored further in the following analyses.

### Region chr1B shows the characteristic features of fungal accessory chromosomes

In many aspects, the region chr1B of *Cd*709 resembled the mini-chromosomes 11 and 12. All three compartments were more AT-rich than the core genome. Region chr1B was also highly enriched with TEs, having 32.8 % coverage with TE copies by length, similar to chr11 and chr12 (32.3 and 35.1 %, respectively), whereas the core chromosomes (excluding chr1B) had only 3 to 6.2 % TE coverage (Table 1, Fig. 3, Table S11). Moreover, the distribution of TE families in region 1B and the two mini-chromosomes differed markedly from the core chromosomes in that they were all enriched with LINE retrotransposons (44 %, 19 % and 34 % coverage, respectively), compared to only 7 % in the core genome (Table 3). LINE TEs are also present in *C. higginsianum* on mini-chromosomes 11 and 12, but their expansion was less striking in this species (7 % and 2 % coverage, respectively) (Fig. S5) than in *Cd*709 [9].

**Table 1:** Characteristics of *C. destructivum* core and mini chromosomes.

|  | *C. destructivum* chromosomes |  |  |  |
| --- | --- | --- | --- | --- |
|  | 1-10<br>(except 1B) | 1B region | 11 | 12 |
| Total length | 48 456 982 bp | 1 205 667 bp | 1 275 594 bp | 812 569 bp |
| G+C content | 54.7 % | 52.3 % | 48.7 % | 50.2 % |
| Number of protein-coding genes | 14882 | 300 | 278 | 171 |
| Proportion of genes by length | 61.7 % | 30.9 %<sup>***</sup> | 32.3 %<sup>***</sup> | 26.8 %<sup>***</sup> |
| Proportion of genes with unknown function | 21.3 % | 42.0 %<sup>***</sup> | 28.4 %<sup>*</sup> | 32.2 %<sup>*</sup> |
| Proportion of genes with RNA support | 77.0 % | 52.0 %<sup>***</sup> | 46.4 %<sup>***</sup> | 59.6 %<sup>*</sup> |
| Proportion of CAZyme genes | 4.1 % | 0.0 %<sup>***</sup> | 1.4 % | 1.2 % |
| Proportion of effector genes | 3.0 % | 4.3 % | 5.4 %<sup>*</sup> | 5.8 %<sup>*</sup> |
| Proportion of SMKG | 0.7 % | 0.0 % | 2.9 %<sup>**</sup> | 0.0 % |
| Proportion of TE by length | 4.4 % | 32.8 %<sup>***</sup> | 32.3 %<sup>***</sup> | 35.1 %<sup>***</sup> |

Asterisks indicate that the data for chromosomes 1B, 11 or 12 differ significantly from the core chromosomes (Fisher's exact test, \*\*\* $P < 0.001$ ; \*\* $P < 0.01$ ; \* $P < 0.05$ )

Examination of the gene content of region chr1B revealed that, similar to the mini-chromosomes, it was overall depleted in protein-coding genes (2-fold less than the core chromosomes), contained a significantly larger proportion of genes encoding proteins of unknown function (i.e. annotated as hypothetical proteins), and had fewer expressed genes (RNA-seq transcript evidence) compared to the core genome (Table 1). Considering categories of potentially pathogenicity-related genes, no CAZyme genes or SMKGs were detected in either region chr1B or chr 12, although eight SMKGs were present on chr11 (Table S6, Tab ‘Secondary metabolism’), all of which had RNA-seq transcript support. Moreover, 38 effectors were found in chr1B and the two mini-chromosomes. Remarkably, 36 of these were absent from *C. higginsianum* (had no RBH in *Ch*63), of which 20 were expressed *in planta* (Table S6 tab ‘Predicted effectors). With 15 and 10 effectors respectively, the mini-chromosomes 11 and 12 were significantly enriched in putative effectors compared to the core chromosomes whereas no enrichment was observed for the 13 effectors of the chr1B (Table 1). Remarkably, the most highly expressed effectors during the biotrophic phase (48 hpi), namely CDEST_01870 (chr1B) and CDEST_15472 (chr12), were located on mini-chromosome-like regions. This raises the possibility that genes carried in such regions are important for virulence.

### Codon usage in region chr1B and the mini-chromosomes differs from the core chromosomes

Analyses of codon usage were used previously to detect differences between the core and accessory chromosomes or lineage-specific compartments of other plant pathogenic fungi [15, 50, 51]. We therefore computed the codon usage of CDS located on the core chromosomes, mini-chromosomes and the chr1B region of *Cd*709. Based on a principal component analysis, codon usage on the core chromosomes was very homogeneous, whereas that of the mini-chromosomes and region chr1B clustered together and separately from the core chromosomes (Fig. 6A). To illustrate this in greater detail, we plotted the codon usage for each amino acid and for each chromosome or region (Fig. S6, representative examples are given for 3 amino acids in Fig. 6B). For these analyses, we excluded the two amino acids (Trp and Met) that are encoded by a single codon. Based on Fisher’s exact tests for each of the remaining 59 codons, almost all the codon usages were different between the core chromosomes on one hand and chr1B, chr11 or chr12 on the other hand. In striking contrast, there were only three differential codon usages between chr1B and chr11 and one between chr1B and chr12. However, chr11 and chr12 were most different from each other with 15 differential codons (Table S12; adjusted *P* < 0.001).

**Figure 6:**
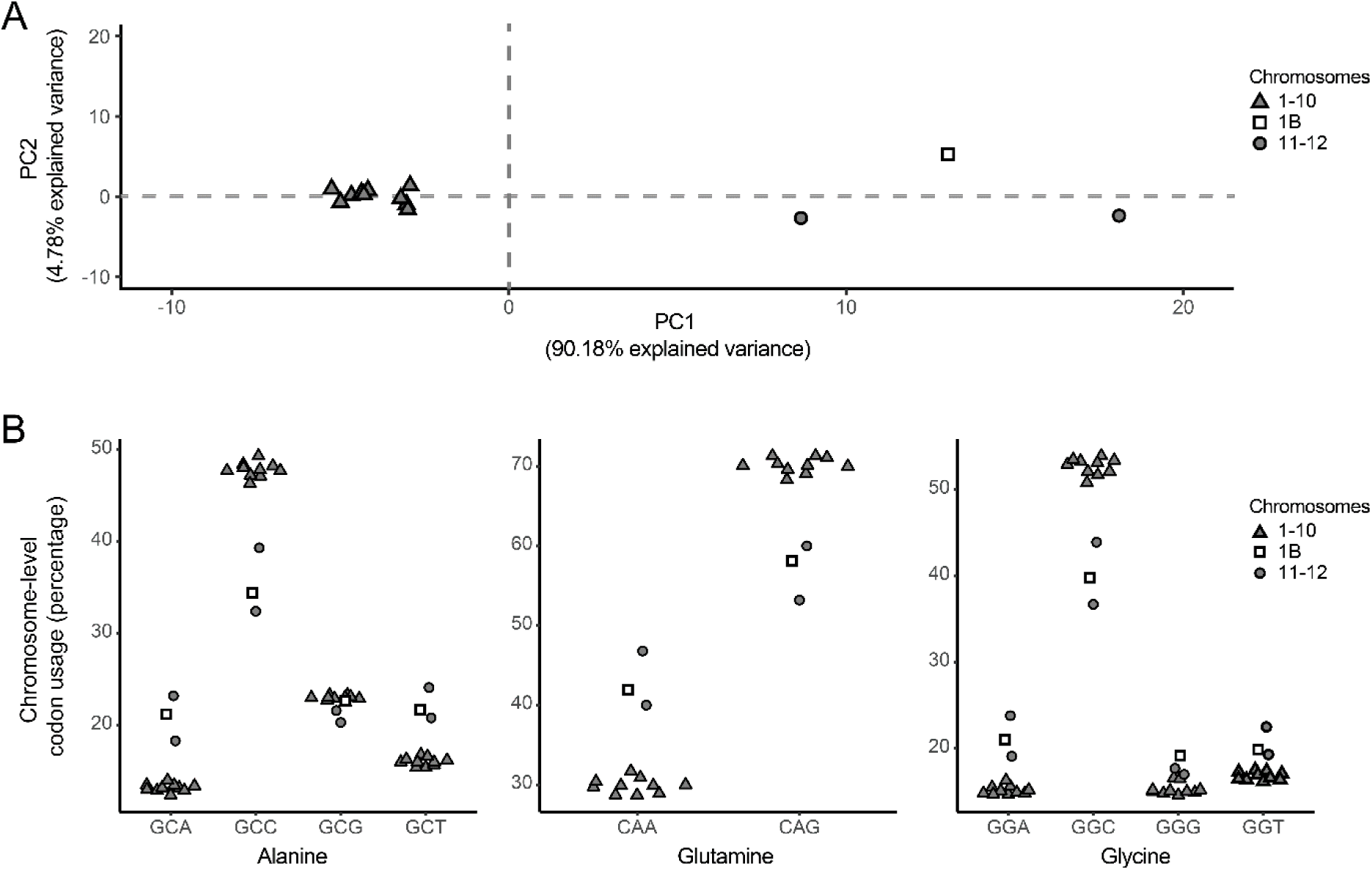
Codon usage bias in the core and mini-chromosomes of *C. destructivum*. **(A)** Principal component analysis (PCA) of codon usage for all amino acids on each chromosome. The region chr1B was considered separately from the rest of chr1. The first two axes accounted for 95% of the variance. **(B)** Plots showing codon usage bias for three amino acids (Alanine, Glutamine, Glycine) in genes located on core chromosomes (1 to 10 excluding region 1B), mini-chromosomes 11 and 12 and region 1B. Codon usage on chr11, chr12 and region chr1B differed significantly from that on core chromosomes (Fisher’s exact test, P < 0.001) for the 10 codons presented except GCG (all comparisons) and GGG (chr12 vs core). Other amino acids are displayed in Fig. S6. The significance is reported for all the codons in the Table S12.

### Region chr1B is a hotspot for segmental duplications

The genome of *Cd*709 was inspected for segmental duplications, as described previously for *C. higginsianum* [9]. A total of 48 duplications involving genes were detected on four chromosomes (chr1, chr6, chr11 and chr12). Among them, 12 duplications were larger than 10 kb (Fig. 7) of which only three were inter-chromosomal (all involving chr12). Similar to *C. higginsianum* [9], these inter-chromosomal duplications were all associated on at least one side with TEs, supporting a potential role of TEs in duplication (Fig. S7). However, in contrast to *C. higginsianum*, these duplications did not take place preferentially near telomeres.

**Figure 7:**
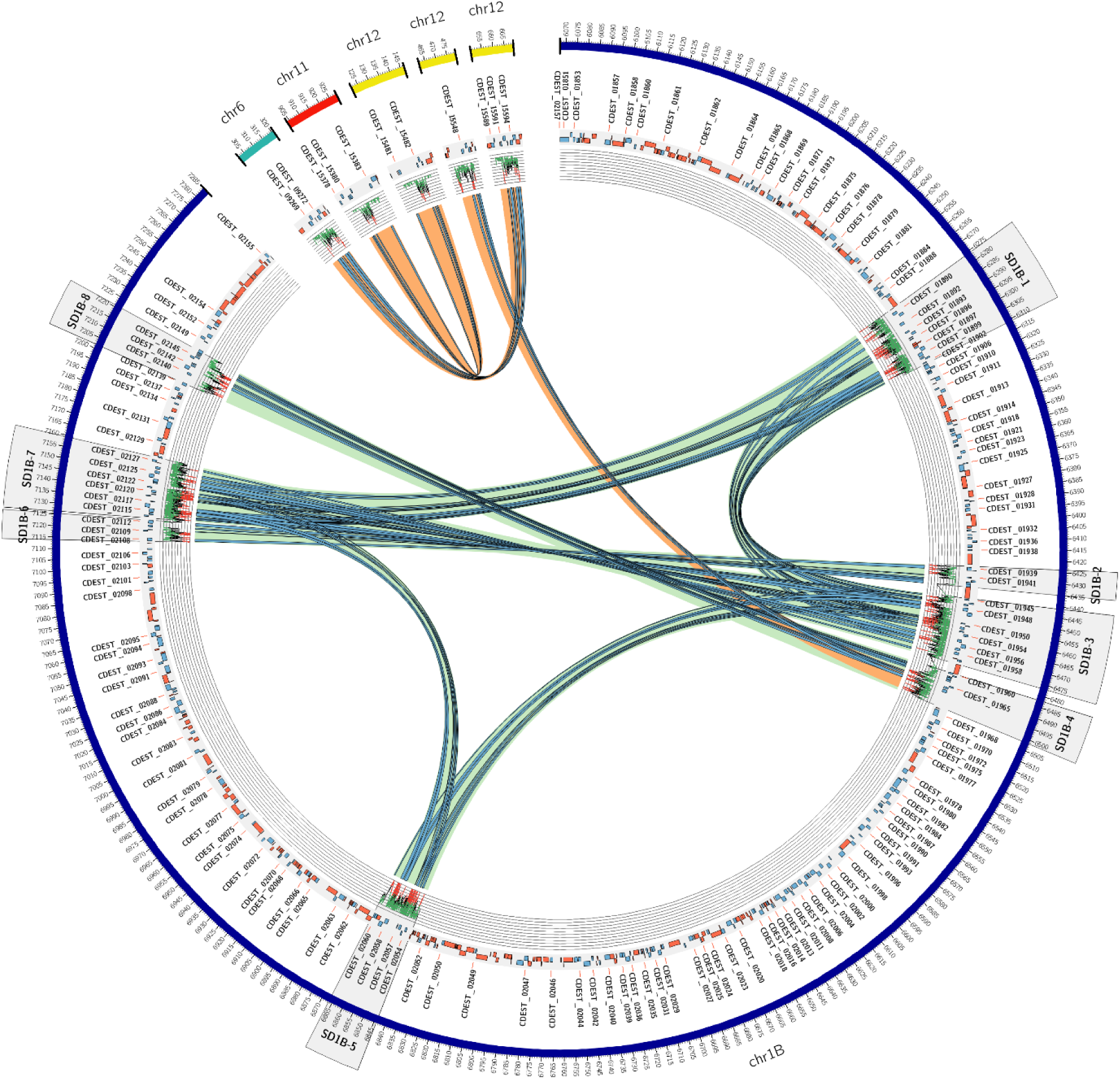
Circos plot showing *Colletotrichum destructivum* segmental duplications larger than 10kb found with SDDetector. The green and red tracks represent genes and transposable elements respectively. The light green and orange arcs indicate intra-chromosomal and inter-chromosomal duplications respectively. Duplicated genes are highlighted by blue arcs. The level of sequence similarity along the duplications is shown by a line graph with a colour scale where green indicates greater than 95% similarity, black between 95 and 90% similarity and red below 90% similarity. A sliding window of 100 base pairs was used to calculate and display sequence similarity from large alignments. The scale displayed on the graph ranges from 100% to 85% similarity.

A remarkable feature of region chr1B was that it showed a strong intra-chromosome duplication pattern, with some regions replicated up to three times (Fig. 7). Assembling large duplications can be difficult even with long-read sequences [52]. To check for possible bias during assembly, the eight largest intra-chromosome duplications on chr1B were inspected manually (Table S13). Due to the problem of multiple reads mapping to duplicated regions, we considered only uniquely mapped reads. Consequently, the read-coverage of these eight regions was on average 2-fold lower than the non-duplicated regions. No other regions of chr1B showed a significant decrease in coverage, and the extremities of the SD regions were well-anchored to chr1B. Reads were identified spanning the two smallest duplications, SD1B-2 (10 reads) and SD1B-6 (22 reads), but other duplicated regions were too large (>16 kb) to be spanned by single PacBio reads. Finally, the short-read RNA-seq data used to annotate the genome were also employed to detect mutations within the duplicated genes. Mutations were detected in all the duplicated regions, albeit with support from only few reads in most cases. Taken together, these results support the reliability of the observed duplications in region chr1B.

To gain insight into the possible origin of region chr1B, we examined conservation of the 300 genes contained within this region in the genomes of 23 other *Colletotrichum* species (Table S14). As expected, given that *C. destructivum* and *C. higginsianum* belong to the same species complex [3], the total proteome of *Cd*709 showed greatest similarity to that of *Ch*63 (14,372 conserved proteins). Surprisingly however, the chr1B proteome shared most conserved proteins with a phylogenetically distant species, namely *C. truncatum* (217 protein matches, compared to only 134 matches in *C. higginsianum*) [53]. Almost half of the genes shared with *C. truncatum* were involved in segmental duplications within the *Cd*709 chr1B. Remarkably, the region triplicated in SD1B-1, SD1B-3 and SD1B-7 was also found in a large duplicated region represented by two contigs within the *C. truncatum* genome assembly (Fig. S8) [54], which may be located on a mini-chromosome due to their low GC content (49.0%, compared to 51.2% in the longer contigs of *C. truncatum*). Other genes located within the *Cd*709 SD1B-1 duplications had Blast matches that were mostly restricted to *C. incanum, C. spaethianum* and *C. tofieldiae* (Spaethianum species complex), *C. salicis* and *C. nymphaeae* (Acutatum species complex), *C. fructicola* (Gloeosporioides species complex), *C. sublineola* (Graminicola species complex) and *C. orchidophilum*, which vary in their phylogenetic distance from *C. destructivum* [53]. The absence of these gene sequences from the *C. higginsianum* genome was confirmed by Tblastn searches against the NCBI wgs Colletotrichum database (266 genomes).

Examination of the gene content in duplicated regions of chr1B gave few clues to their possible role in the host interaction or the advantage for the fungus to maintain multiple mutated copies of these genes. One gene duplicated four times (CDEST_01898, CDEST_01949, CDEST_02058 and CDEST_02116) encoded a major facilitator superfamily transporter. The five genes duplicated between SD1B-2 and SD1B-6 comprised four FAD-binding domain-containing proteins and a patatin-like serine hydrolase.

## Discussion

In this study, we present a chromosome-level reference assembly of the *C. destructivum* genome, a phytopathogen causing anthracnose disease principally on species of *Medicago* and *Trifolium* (Fabaceae). Among other members of the Destructivum species complex, which currently contains 17 recognised species [3], the genomes of *C. lentis, C. tanaceti* and *C. shisoi* were sequenced previously but the resulting assemblies were highly fragmented, containing 2980, 5242 and 36,350 contigs, respectively [8, 11, 12]. Using PacBio long-read sequencing, we were able to generate a gapless assembly of the *Cd*709 genome which, together with that of *Ch*63 [9], provides a second complete genome within the Destructivum species complex, facilitating future comparative genomic analyses within this important group of plant pathogens.

Alignment of the *Cd*709 genome assembly with those of *C. higginsianum* strains *Ch*63 and *Ch*35 revealed large-scale chromosome rearrangements between the two closely-related species. Some of these rearrangements were potentially mediated by recombination between homologous regions containing TEs, which flanked one or both of the breakpoints. Similar TE-mediated chromosome rearrangements were previously reported at the intra-species level in *C. higginsianum* [10]. Our analysis of synteny between the genomes of *Cd*709 and *Ch*63 also revealed the presence of a 1.2 Mb species-specific region within Chr1 of *Cd*709, which we called Chr1B. This ‘accessory region’ (AR) displays many of the hallmarks that characterize fungal mini-chromosomes, or ‘accessory chromosomes’, in that it is AT-rich, transposon-rich, gene-poor and has a distinct codon usage [51, 55–57]. In all these respects, Chr1B resembles the mini-chromosomes Chr11 and Chr12 but is strikingly different from the rest of Chr1 and other core chromosomes of *Cd*709. The TE enrichment observed in Chr1B and both mini-chromosomes is largely caused by the specific expansion of LINE and TIR elements in these compartments, unlike the core chromosomes where the Gipsy TE family predominates.

Using PFGE and Southern hybridization with a probe specific to Chr1B, we were able to confirm that this AR is carried not only on Chr1 of *Cd*709 but also on the largest chromosome of *Cd*202, despite the widely-separated geographical origins of these two isolates (Saudi Arabia and Morocco, respectively). Analysis of a larger collection of *C. destructivum* isolates is now needed to determine the extent to which Chr1B is conserved within this pathogen species. The presence of an AR embedded within a core chromosome has been reported in other plant pathogenic fungi. For example, isolates of the T race of *Cochliobolus heterostrophus* harbor an AR of about 1.2 Mb distributed between two core chromosomes that contains the *Tox1* locus producing the T-toxin polyketide [58, 59]. In *Verticillium dahliae*, Chr3 and Chr4 each harbor two ARs of ∼300 kb [60], while in *Fusarium poae* a 204 kb block with AR characteristics is inserted near one telomere of Chr3 [57]. However, it should be noted that in these two examples the inserted AR blocks are 4- to 6-fold smaller than Chr1B of *Cd*709.

Our working hypothesis is that the AR Chr1B arose by the integration of a mini-chromosome into a core chromosome of *C. destructivum*, but the mechanism by which this occurred is unclear. Despite the subtelomeric position of Chr1B, its integration is unlikely to have resulted from the telomeric fusion of a mini-chromosome with a core chromosome because it is flanked on both sides by portions of Chr1, both of which are highly syntenic to Chr2 of *C. higginsianum*. A chromosome containing distinct regions characteristic of core and accessory chromosomes was previously reported in the genome of *C. fructicola* strain Nara gc5 [61]. In this case, the chimeric chromosome, called Nara_c11, is smaller (2.8 Mb) than *Cd*709 chromosome 1 (7.3 Mb) and the TE-rich, gene-poor AR occupies most of the chromosome (66 % by length), in contrast to *Cd*709 Chr1B, which occupies only 16 %. A further difference to *Cd*709 Chr1B is that the AR of Nara_c11 includes a telomere, suggesting that in this case the chimeric chromosome arose through a different mechanism. Taken together, our findings provide further evidence for genetic exchange between core and accessory genomic compartments in *Colletotrichum* species [61]. In other fungi, chromosome breakage-fusion-bridge (BFB) cycles have been invoked not only in the creation of accessory chromosomes from core chromosomes [62], but also in their reintegration into core chromosomes [63].

A distinguishing feature of the Chr1B AR is that it has undergone extensive region-specific segmental duplications. Some inter-chromosomal SDs in *Cd*709 were associated with TEs at one or both of their borders, as we found previously in *Ch*63 [9], but there was little evidence that the region-specific SDs in Chr1B were mediated by TEs. Similarly, the AR of *C. fructicola* chromosome Nara_c11 was found to be implicated in numerous intra- and inter-chromosomal SDs but as in *Cd*709 these were not consistently flanked by TEs [61]. Among fungal pathogens, SDs can play important roles in generating genetic diversity and novel gene functions, either at the level of expression or coding sequence [64, 65]. A recent study on *Fusarium* strains infecting banana also highlighted the importance of SDs in driving the evolution of ARs and the effector genes contained within them [66]. Although the *C. destructivum* genome contains a complete Mat1-2-1 mating-type locus (Table S6, Tab MAT1-2-1), and should therefore be capable of sexual reproduction, this has never been observed [67], [3]. In this context, segmental duplication may therefore provide an important mechanism for generating genetic diversity for host adaptation in this essentially asexual pathogen.

A remarkable finding was that some segmentally duplicated blocks of genes within Chr1B of *C. destructivum* are conserved and syntenic with duplicated regions in the genome of *C. truncatum*, a species that is phylogenetically very distant [53]. Given that these two taxa diverged ∼60 million years ago [68], soon after speciation in *Colletotrichum*, these SDs may be very ancient and have been selectively retained in some species and lost in others. Alternatively, these duplicated regions may have been acquired by horizontal chromosome transfer (HCT) from another species to a common ancestor, or through independent transfers to *C. destructivum* and *C. truncatum*. HCT would be consistent with the distinct codon bias in Chr1B and the taxonomic incongruity of many genes within this region. The horizontal transfer of a mini-chromosome between vegetatively incompatible biotypes of *C. gloeosporioides* was shown experimentally [69, 70], and it is well-documented that genetic material can be exchanged following fusion between conidial anastomosis tubes of the same, or even different, *Colletotrichum* species [71–73].

Chr1B contains a variety of genes with potential roles in fungal virulence, some of which were expressed during infection. These include genes encoding 13 candidate secreted effector proteins, 8 protein kinases, 5 major facilitator superfamily membrane transporters, 5 heterokaryon incompatibility (HET) proteins and 8 putative transcription factors (TFs) (Table S6). It is interesting to note that, similar to Chr1B, the accessory ‘pathogenicity chromosome’ of *Fusarium oxysporum* f.sp*. lycopersici* is enriched not only with effectors genes but also with genes encoding protein kinases, membrane transporters, HET proteins and TFs, of which one TF was shown to regulate the expression of plant-induced effector genes [74],[75]. TFs were also found to be enriched in the four lineage-specific ARs of *V. dahliae* [60]. Overall, the gene content of Chr1B suggests that it may contribute to *C. destructivum* pathogenicity. This was demonstrated experimentally for ARs in two other members of the Destructivum species complex, namely Chr11 of *C. higginsianum* (isolate *Ch*35) which was essential for virulence on *A. thaliana* [17], and Chr11 of *C. lentis*, which was required for virulence on lentil [12]. In the case of *Cd*709, it is noteworthy that the three most highly expressed and plant-induced effector genes are all located in ARs, namely CDEST_01870 on Chr1B, CDEST_15404 on Chr11 and CDEST_15472 on Chr12. These and other pathogenicity-related genes carried within these genomic compartments will provide interesting candidates for future functional analysis.

Finally, we show here that *Cd*709 can complete its life cycle not only on its original host, *M. sativa*, but also on the widely-studied model legume, *M. truncatula*. Until now, the only other *Colletotrichum* species known to attack *M. truncatula* was *C. trifolii*, which belongs to the phylogenetically distant Orbiculare species complex and uses a different infection process where the biotrophic phase extends to many host cells [76, 77]. With complete genome assemblies and high-quality gene annotations available for both partners, together with abundant genetic tools and resources on the plant side, the *C. destructivum* - *M. truncatula* interaction could provide a tractable new model pathosystem for studying hemibiotrophic fungal interactions with Fabaceae hosts. Our identification of susceptible and resistant *M. truncatula* accessions also raises the possibility that natural variation among accessions could be exploited to analyse the genetic basis of resistance to *C. destructivum* [78].

## Author contributions

Conceptualization: NL, AS, CK, JFD, RJO; Investigation: PLP, SP, AA, RJO; Formal analysis: NL, AS, AL, JA, JFD; Visualization: NL, AS, PLP, JA, CK, JFD, RJO; Writing – Original Draft: NL, AS, PLP, JA, CK, JFD, RJO; Writing – Review & Editing: NL, AS, CK, JFD, RJO; Supervision: NL, CK, RJO; Funding acquisition: RJO.

## Acknowledgements

We are grateful to the following bioinformatics platforms and partners for providing help and/or computing and/or storage resources: Genotoul bioinformatics platform Toulouse Occitanie (Bioinfo Genotoul, doi: 10.15454/1.5572369328961167E12), CATI BARIC (https://www.cesgo.org/catibaric/), and INRAE-LIPME Bioinfo (Sébastien Carrère).We also thank Dr Alexander Wittenberg (KeyGene N.V.,The Netherlands) for help with sequencing, and the INRAE Centre de Ressources Biologiques *Medicago truncatula* for providing seeds.

## Abbreviations

AT: acyltransferase domain
AR: accessory region
BDBH: bidirectional best hit
BGC: biosynthetic genes cluster
CAT: conidial anastomosis tubes
CAZyme: carbohydrate active enzyme
CDS: coding sequence
CE: carbohydrate esterase
Chr: chromosome
DNA: deoxyribonucleic acid
GH: glycoside hydrolase
GO: gene ontology
HCT: horizontal chromosome transfer
HPI: hours post-inoculation
KS: ketosynthase domain
LINE: long interspersed nuclear element
LTR: long terminal repeats
MITE: miniature inverted-repeat transposable element
NRPS: non-ribosomal peptide synthetase
PCA: principal component analysis
PCP: peptidyl carrier protein domain
PCR: polymerase chain reaction
PFGE: pulsed-field gel electrophoresis
PKS: polyketide synthase
PL: polysaccharide lyase
RBH: reciprocal best hit
RFP: red fluorescent protein
RNA: ribonucleic acid
SD: segmental duplication
SMKG: secondary metabolism key gene
SMRT: single molecule real time
TE: transposable element
TIR: terminal inverted repeat
TPM: transcript per million.

## References

1. Frayssinet S. *Colletotrichum destructivum*: a new lucerne pathogen in Argentina. Australasian Plant Disease Notes 2008 3:1 2008;3:68–68.

2. Latunde-Dada AO, Bailey JA, Lucas JA. Infection process of *Colletotrichum destructivum* O’Gara from lucerne (Medicago sativa L.). European Journal of Plant Pathology 1997;103:35–41.

3. Damm U, O’Connell RJ, Groenewald JZ, Crous PW. The *Colletotrichum destructivum* species complex – hemibiotrophic pathogens of forage and field crops. Studies in Mycology 2014;79:49–84.

4. Sun HY, Liang Y. First report of anthracnose on sunflower caused by *Colletotrichum destructivum* in China. Plant Disease 2018;102:245.

5. Manandhar JB. *Colletotrichum destructivum*, the anamorph of *Glomerella glycines*. Phytopathology 1986;76:282.

6. Tiffany LH, Gilman JC. Species of *Colletotrichum* from Legumes. Mycologia 1954;46:52–75.

7. Damm U, Sato T, Alizadeh A, Groenewald JZ, Crous P. The *Colletotrichum dracaenophilum*, *C. magnum* and *C. orchidearum* species complexes. Studies in mycology;92. 2019. DOI: 10.1016/J.SIMYCO.2018.04.001.

8. Gan P, Tsushima A, Hiroyama R, Narusaka M, Takano Y, et al. *Colletotrichum shisoi* sp. nov., an anthracnose pathogen of *Perilla frutescens* in Japan: molecular phylogenetic, morphological and genomic evidence. Scientific Reports;9. 2019. DOI: 10.1038/s41598-019-50076-5.

9. Dallery J-F, Lapalu N, Zampounis A, Pigné S, Luyten I, et al. Gapless genome assembly of *Colletotrichum higginsianum* reveals chromosome structure and association of transposable elements with secondary metabolite gene clusters. BMC Genomics 2017;18:667.

10. Tsushima A, Gan P, Kumakura N, Narusaka M, Takano Y, et al. Genomic plasticity mediated by transposable elements in the plant pathogenic fungus *Colletotrichum higginsianum*. Genome biology and evolution 2019;11:1487–1500.

11. Lelwala R V., Korhonen PK, Young ND, Scott JB, Ades PK, et al. Comparative genome analysis indicates high evolutionary potential of pathogenicity genes in *Colletotrichum tanaceti*. PLOS ONE 2019;14:e0212248.

12. Bhadauria V, MacLachlan R, Pozniak C, Cohen-Skalie A, Li L, et al. Genetic map-guided genome assembly reveals a virulence-governing minichromosome in the lentil anthracnose pathogen *Colletotrichum lentis*. The New phytologist 2019;221:431–445.

13. Wang Haoming, Huang Rong, Ren Jingyi, Tang Lihua, Huang Suiping, et al. The evolution of mini-chromosomes in the fungal genus *Colletotrichum*. mBio 2023;14:e00629–23.

14. Plaumann P-L, Koch C. The many questions about mini chromosomes in *Colletotrichum spp*. Plants;9. 2020. DOI: 10.3390/plants9050641.

15. Ma LJ, Van Der Does HC, Borkovich KA, Coleman JJ, Daboussi MJ, et al. Comparative genomics reveals mobile pathogenicity chromosomes in *Fusarium*. Nature 2010 464:7287 2010;464:367–373.

16. Liu S, Lin G, Ramachandran SR, Daza LC, Cruppe G, et al. Rapid mini-chromosome divergence among fungal isolates causing wheat blast outbreaks in Bangladesh and Zambia. New Phytologist. 2023. DOI: 10.1111/nph.19402.

17. Plaumann P-L, Schmidpeter J, Dahl M, Taher L, Koch C. A dispensable chromosome Is required for virulence in the hemibiotrophic plant pathogen *Colletotrichum higginsianum*. Frontiers in microbiology 2018;9:1005.

18. O’Connell R, Herbert C, Sreenivasaprasad S, Khatib M, Esquerré-Tugayé M-T, et al. A novel *Arabidopsis-Colletotrichum* pathosystem for the molecular dissection of plant-fungal interactions. Molecular plant-microbe interactions : MPMI 2004;17:272–82.

19. Stiehler F, Steinborn M, Scholz S, Dey D, Weber APM, et al. Helixer: cross-species gene annotation of large eukaryotic genomes using deep learning. Bioinformatics 2021;36:5291–5298.

20. Koren S, Walenz BP, Berlin K, Miller JR, Bergman NH, et al. Canu: scalable and accurate long-read assembly via adaptive k-mer weighting and repeat separation. Genome research 2017;27:722–736.

21. Seppey M, Manni M, Zdobnov EM. BUSCO: Assessing genome assembly and annotation completeness. In: Methods in Molecular Biology. Humana Press Inc.; 2019. pp. 227–245.

22. Rehmeyer C, Li W, Kusaba M, Kim Y-S, Brown D, et al. Organization of chromosome ends in the rice blast fungus, *Magnaporthe oryzae*. Nucleic Acids Research 2006;34:4685–4701.

23. Soorni A, Haak D, Zaitlin D, Bombarely A. Organelle_PBA, a pipeline for assembling chloroplast and mitochondrial genomes from PacBio DNA sequencing data. BMC genomics 2017;18:49.

24. Bolger AM, Lohse M, Usadel B. Trimmomatic: a flexible trimmer for Illumina sequence data. Bioinformatics (Oxford, England) 2014;30:2114–20.

25. Dobin A, Davis CA, Schlesinger F, Drenkow J, Zaleski C, et al. STAR: ultrafast universal RNA-seq aligner. Bioinformatics 2013;29:15–21.

26. Flutre T, Duprat E, Feuillet C, Quesneville H. Considering transposable element diversification in *de novo* annotation approaches. PloS one 2011;6:e16526.

27. Amselem J, Lebrun M-H, Quesneville H. Whole genome comparative analysis of transposable elements provides new insight into mechanisms of their inactivation in fungal genomes. BMC genomics 2015;16:141.

28. Hoede C, Arnoux S, Moisset M, Chaumier T, Inizan O, et al. PASTEC: an automatic transposable element classification tool. PLoS ONE 2014;9:e91929.

29. Wicker T, Sabot F, Hua-Van A, Bennetzen JL, Capy P, et al. A unified classification system for eukaryotic transposable elements. Nature Reviews Genetics 2007;8:973–982.

30. Sallet E, Gouzy J, Schiex T. EuGene: An automated integrative gene finder for eukaryotes and prokaryotes. Humana, New York, NY; 2019. pp. 97–120.

31. Min B, Grigoriev I V, Choi I-G. FunGAP: fungal genome annotation pipeline using evidence-based gene model evaluation. Bioinformatics 2017;33:2936–2937.

32. Eilbeck K, Moore B, Holt C, Yandell M. Quantitative measures for the management and comparison of annotated genomes. BMC bioinformatics 2009;10:67.

33. Valach M, Burger G, Gray MW, Lang BF. Widespread occurrence of organelle genome-encoded 5S rRNAs including permuted molecules. Nucleic acids research 2014;42:13764–77.

34. Bernt M, Donath A, Jühling F, Externbrink F, Florentz C, et al. MITOS: improved *de novo* metazoan mitochondrial genome annotation. Molecular Phylogenetics and Evolution 2013;69:313–319.

35. Drillon G, Carbone A, Fischer G. SynChro: a fast and easy tool to reconstruct and visualize synteny blocks along eukaryotic chromosomes. PLoS ONE 2014;9:e92621.

36. Gilchrist CLM, Chooi YH. clinker & clustermap.js: automatic generation of gene cluster comparison figures. Bioinformatics (Oxford, England) 2021;37:2473–2475.

37. Jones P, Binns D, Chang H-Y, Fraser M, Li W, et al. InterProScan 5: genome-scale protein function classification. Bioinformatics 2014;30:1236.

38. Camacho C, Coulouris G, Avagyan V, Ma N, Papadopoulos J, et al. BLAST+: architecture and applications. BMC Bioinformatics 2009;10:421.

39. Gene Ontology Consortium. The Gene Ontology (GO) database and informatics resource. Nucleic Acids Research 2004;32:258D –261.

40. Götz S, García-Gómez JM, Terol J, Williams TD, Nagaraj SH, et al. High-throughput functional annotation and data mining with the Blast2GO suite. Nucleic Acids Research 2008;36:3420.

41. Zhang H, Yohe T, Huang L, Entwistle S, Wu P, et al. dbCAN2: a meta server for automated carbohydrate-active enzyme annotation. Nucleic Acids Research 2018;46:W95–W101.

42. Petersen TN, Brunak S, von Heijne G, Nielsen H. SignalP 4.0: discriminating signal peptides from transmembrane regions. Nature Methods 2011;8:785–786.

43. Emanuelsson O, Nielsen H, Brunak S, von Heijne G. Predicting subcellular localization of proteins based on their N-terminal amino acid sequence. Journal of Molecular Biology 2000;300:1005–1016.

44. Krogh A, Larsson B, von Heijne G, Sonnhammer ELL. Predicting transmembrane protein topology with a hidden markov model: application to complete genomes. Journal of Molecular Biology 2001;305:567–580.

45. Sperschneider J, Dodds PN, Gardiner DM, Singh KB, Taylor JM. Improved prediction of fungal effector proteins from secretomes with EffectorP 2.0. Molecular plant pathology 2018;19:2094–2110.

46. Blin K, Shaw S, Steinke K, Villebro R, Ziemert N, et al. antiSMASH 5.0: updates to the secondary metabolite genome mining pipeline. Nucleic Acids Research 2019;47:W81–W87.

47. Zhou P, Silverstein KAT, Ramaraj T, Guhlin J, Denny R, et al. Exploring structural variation and gene family architecture with *de novo* assemblies of 15 Medicago genomes. BMC Genomics 2017;18:261.

48. Moll KM, Zhou P, Ramaraj T, Fajardo D, Devitt NP, et al. Strategies for optimizing BioNano and Dovetail explored through a second reference quality assembly for the legume model, *Medicago truncatula*. BMC Genomics 2017;18:578.

49. O’Connell RJ, Thon MR, Hacquard S, Amyotte SG, Kleemann J, et al. Lifestyle transitions in plant pathogenic *Colletotrichum* fungi deciphered by genome and transcriptome analyses. Nature Genetics 2012;44:1060–1065.

50. Coleman JJ, Rounsley SD, Rodriguez-Carres M, Kuo A, Wasmann CC, et al. The genome of *Nectria haematococca*: contribution of supernumerary chromosomes to gene expansion. PLOS Genetics 2009;5:e1000618.

51. Hu J, Chen C, Peever T, Dang H, Lawrence C, et al. Genomic characterization of the conditionally dispensable chromosome in *Alternaria arborescens* provides evidence for horizontal gene transfer. BMC genomics 2012;13:1–13.

52. Vollger MR, Dishuck PC, Sorensen M, Welch AE, Dang V, et al. Long-read sequence and assembly of segmental duplications. Nature Methods 2019;16:88–94.

53. Liu F, Ma ZY, Hou LW, Diao YZ, Wu WP, et al. Updating species diversity of *Colletotrichum*, with a phylogenomic overview. Studies in Mycology;101.

54. Rogério F, Boufleur TR, Ciampi-Guillardi M, Sukno SA, Thon MR, et al. Genome sequence resources of *Colletotrichum truncatum*, *C. plurivorum*, C. musicola, and C. sojae: four species pathogenic to soybean (Glycine max). Phytopathology 2020;110:1497–1499.

55. Langner T, Harant A, Gomez-Luciano LB, Shrestha RK, Win J, et al. Genomic rearrangements generate hypervariable mini-chromosomes in host-specific lineages of the blast fungus. bioRxiv 2020;2020.01.10.901983.

56. Langner T, Harant A, Gomez-Luciano LB, Shrestha RK, Malmgren A, et al. Genomic rearrangements generate hypervariable mini-chromosomes in host-specific isolates of the blast fungus. PLOS Genetics 2021;17:e1009386.

57. Vanheule A, Audenaert K, Warris S, van de Geest H, Schijlen E, et al. Living apart together: crosstalk between the core and supernumerary genomes in a fungal plant pathogen. BMC Genomics 2016;17:1–18.

58. Condon BJ, Leng Y, Wu D, Bushley KE, Ohm RA, et al. Comparative genome structure, secondary metabolite, and effector coding capacity across *Cochliobolus* pathogens. PLoS Genetics 2013;9:e1003233.

59. Yang G, Turgeon BG, Yoder OC, Bronson CR, Yoder OC, et al. Toxin-deficient mutants from a toxin-sensitive transformant of *Cochliobolus heterostrophus*. Genetics 1994;137:751–7.

60. Klosterman SJ, Subbarao K V., Kang S, Veronese P, Gold SE, et al. Comparative genomics yields insights into niche adaptation of plant vascular wilt pathogens. PLOS Pathogens 2011;7:e1002137.

61. Gan P, Hiroyama R, Tsushima A, Masuda S, Shibata A, et al. Telomeres and a repeat-rich chromosome encode effector gene clusters in plant pathogenic *Colletotrichum* fungi. Environmental Microbiology 2021;23:6004–6018.

62. Croll D, Zala M, McDonald BA. Breakage-fusion-bridge cycles and large insertions contribute to the rapid evolution of accessory chromosomes in a fungal pathogen. PLOS Genetics 2013;9:e1003567.

63. Bertazzoni S, Williams AH, Jones DA, Syme RA, Tan K-C, et al. Accessories make the outfit: accessory chromosomes and other dispensable DNA regions in plant-pathogenic fungi. Molecular Plant-Microbe Interactions 2018;31:779–788.

64. Francis A, Ghosh S, Tyagi K, Prakasam V, Rani M, et al. Evolution of pathogenicity-associated genes in *Rhizoctonia solani* AG1-IA by genome duplication and transposon-mediated gene function alterations. BMC Biology 2023;21:1–19.

65. Fraser JA, Huang JC, Pukkila-Worley R, Alspaugh JA, Mitchell TG, et al. Chromosomal translocation and segmental duplication in *Cryptococcus neoformans*. Eukaryotic Cell 2005;4:401–406.

66. van Westerhoven A, Aguilera-Galvez C, Nakasato-Tagami G, Shi-Kunne X, Martinez E, et al. Segmental duplications drive the evolution of accessory regions in a major crop pathogen. bioRxiv 2023;2023.06.07.544053.

67. Wilson AM, Lelwala R V., Taylor PWJ, Wingfield MJ, Wingfield BD. Unique patterns of mating pheromone presence and absence could result in the ambiguous sexual behaviors of *Colletotrichum* species. G3 Genes|Genomes|Genetics;11. 2021. DOI: 10.1093/G3JOURNAL/JKAB187.

68. Bhunjun CS, Phukhamsakda C, Jeewon R, Promputtha I, Hyde KD. Integrating different lines of evidence to establish a novel Ascomycete genus and family (*Anastomitrabeculia*, Anastomitrabeculiaceae) in Pleosporales. Journal of Fungi 2021, Vol 7, Page 94 2021;7:94.

69. Manners JM, He C. Slow-growing heterokaryons as potential intermediates in supernumerary chromosome transfer between biotypes of *Colletotrichum gloeosporioides*. Mycological Progress 2011;10:383–388.

70. He C, Rusu AG, Poplawski AM, Irwin JAG, Manners JM. Transfer of a supernumerary chromosome between vegetatively incompatible biotypes of the fungus *Colletotrichum gloeosporioides*. Genetics 1998;150:1459–1466.

71. Roca MG, Davide LC, Davide LMC, Mendes-Costa MC, Schwan RF, et al. Conidial anastomosis fusion between *Colletotrichum* species. Mycological research 2004;108:1320–1326.

72. Ishikawa FH, Souza EA, Shoji JY, Connolly L, Freitag M, et al. Heterokaryon incompatibility is suppressed following conidial anastomosis tube fusion in a fungal plant pathogen. PloS one;7. 2012. DOI: 10.1371/JOURNAL.PONE.0031175.

73. Mehta N, Baghela A. Quorum sensing-mediated inter-specific conidial anastomosis tube fusion between *Colletotrichum gloeosporioides* and *C. siamense*. IMA fungus;12. 2021. DOI: 10.1186/S43008-021-00058-Y.

74. Schmidt SM, Houterman PM, Schreiver I, Ma L, Amyotte S, et al. MITEs in the promoters of effector genes allow prediction of novel virulence genes in *Fusarium oxysporum*. BMC genomics 2013;14:1–21.

75. van der Does HC, Fokkens L, Yang A, Schmidt SM, Langereis L, et al. Transcription factors encoded on core and accessory chromosomes of *Fusarium oxysporum* induce expression of effector genes. PLOS Genetics 2016;12:e1006401.

76. Mould MJR, Boland GJ, Robb J. Ultrastructure of the *Colletotrichum trifolii-Medicago sativa* pathosystem. I. Pre-penetration events. Physiological and Molecular Plant Pathology 1991;38:179–194.

77. Damm U, Cannon PF, Liu F, Barreto RW, Guatimosim E, et al. The *Colletotrichum orbiculare* species complex: Important pathogens of field crops and weeds. Fungal Diversity 2013;61:29–59.

78. C A-T, BB W, MS O, S D, H Z, et al. Identification and characterization of nucleotide-binding site-leucine-rich repeat genes in the model plant *Medicago truncatula*. Plant physiology;146. 2008. DOI: 10.1104/PP.107.104588.

